# Thermal and Chemical Unfolding of a recombinant monoclonal IgG1 antibody: Application of the Multi-State Zimm-Bragg Theory

**DOI:** 10.1101/692236

**Authors:** P. Garidel, A. Eiperle, M. Blech, J. Seelig

## Abstract

The thermal unfolding of a recombinant monoclonal antibody IgG1 (mAb) was measured with differential scanning calorimetry (DSC). The DSC thermograms reveal a pre-transition at 72°C with an unfolding enthalpy of ΔH_cal_ ∼ 200-300 kcal/mol and a main transition at 85 °C with an enthalpy of ∼900 - 1000 kcal/mol. In contrast to single-domain molecules, mAb unfolding is a complex reaction that is analysed with the multi-state Zimm-Bragg theory. For the investigated mAb, unfolding is characterised by a cooperativity parameter σ ∼10^−4^ and a Gibbs free energy of unfolding of g_nu_ ∼100 cal/mol per amino acid. The enthalpy of unfolding provides the number of amino acid residues *v* participating in the unfolding reaction. On average, *v*∼220±50 amino acids are involved in the pre-transition and *v*∼850±30 in the main transition, accounting for ∼90% of all amino acids. Thermal unfolding was further studied in the presence of guanidineHCl. The chemical denaturant reduces the unfolding enthalpy ΔH_cal_ and lowers the midpoint temperature T_0_. Both parameters depend linearly on the concentration of denaturant. The guanidineHCl concentrations needed to unfold mAb at 25 °C are predicted to be 2-3 M for the pre-transition and 5-7 M for the main transition, varying with pH. GuanidineHCl binds to mAb with an exothermic binding enthalpy, which partially compensates the endothermic mAb unfolding enthalpy. The number of guanidineHCL molecules bound upon unfolding is deduced from the DSC thermograms. The bound guanidineHCl-to-unfolded amino acid ratio is 0.79 for the pre-transition and 0.55 for the main transition. The pre-transition binds more denaturant molecules and is more easily destabilised than the main transition.

Overall, the current study shows the strength of the Zimm-Bragg model for the quantitative description of unfolding events of large, therapeutic proteins, such as a monoclonal antibody.

**Statement of significance:** First quantitative thermodynamic study of an antibody with differential scanning calorimetry and analyzed with the multi-state Zimm-Bragg theory.

## Introduction

The standard unfolding model for small proteins (e.g. single-domain molecules) is the 2-state model. Only two types of molecules exist in solution, the native protein (N) and its structural unfolded conformation (U) (all-or-none model) (1). However, “peptides that form helices in solution do not show a simple 2-state equilibrium between a fully folded and a fully unfolded structure. Instead they form a complex mixture of all helix, all coil or, most frequently central helices with frayed coil ends” (2). A more realistic model is the multi-state Zimm-Bragg theory, originally developed for the temperature-induced coil-to-α-helix transition (3; 4). It has been applied successfully to describe the cooperative thermal unfolding of a variety of proteins (5). In fact, the Zimm-Bragg theory provides a perfect quantitative description of the thermal unfolding of apolipoprotein A-1, a protein with a high α-helix content (∼50%) (6; 7) as well as for a large number of other globular proteins (5). In this study, the theory is extended to the unfolding of a large multi-domain protein, a monoclonal antibody of molecular weight 143 kDa where the 2-state model fails completely. The thermal unfolding of a monoclonal antibody mAb was investigated with differential scanning calorimetry (DSC) in buffer and in the presence of the chemical denaturant guanidineHCl. GuanidineHCl is one of the most commonly used chemicals to induce protein unfolding. Increasing the concentration of denaturant shifts the folding equilibrium towards the unfolded state. The molecular mechanism of chemical denaturation is still discussed controversially (8). One theory postulates an indirect mechanism by which chemical denaturants change the water structure and thereby reduce the magnitude of the hydrophobic effect. The alternative view is a direct interaction of the denaturant with the protein (9; 10). Strong support for this mechanism comes from isothermal titration calorimetry (ITC) which provides evidence for an exothermic binding reaction of guanidineHCl with proteins (11). Molecular dynamics simulations (12) and X-ray studies (13) also support a direct interaction mechanism.

In the present study, we used DSC to investigate the stability of mAb as a function of temperature, solvent pH, and guanidineHCl concentration. The thermograms were analysed with the multi-state Zimm-Bragg theory. The investigated mAb exhibits two transitions, a first transition denoted as pre-transition and a second transition at higher temperature connected to a higher transition enthalpy denoted as main transition. These transitions are characterised by their enthalpy, midpoint transition temperature, and molar heat capacity. The transitions result from the denaturation of specific domains of the monoclonal antibody. For the investigated IgG1, the first transition is probably related to the reversible transition of C_H_2 domains whereas the large, irreversible transition at high temperature is tentatively assigned to Fab and C_H_3 domains (14; 15).

GuanidineHCl was added up to a concentration of 2.5 M. The chemical denaturant destabilized the antibody, decreasing midpoint temperatures T_0_ and unfolding enthalpy ΔH_cal_. The molecular mechanism behind destabilization is deduced from the unfolding enthalpy.

## Materials and Methods

### mAb sample preparation

The humanized recombinant monoclonal antibody (mAb) of IgG1 isotype was produced by mammalian cell culture technology and purified accordingly (16; 17). The concentration of the IgG1 sample solutions were determined by UV-measurement at 280 nm using an extinction coefficient of 1.32 for a 1 mg/ml solution (path length d = 1 cm).

Purity was determined by size exclusion chromatography. The monomer content as measured by HPSEC was >99% (18).

The pH of the sample was varied by titrating HCl, respective NaOH, to obtain the target pH value as described in the text. GuanidineHCl was added to the protein sample to generate a concentration range from 0 to 2.5 M. If necessary, the pH was readjusted after the addition of guanidineHCl.

### Analysis of mAb differential scanning calorimetry (DSC) thermograms

Protein concentrations were typically 3 mg/mL corresponding to ∼20 μM. Starting at 5 °C the thermal unfolding of mAb was measured by increasing the temperature to 95 °C at a heating scan rate of 1 K/min. DSC experiments were performed with a VP-DSC instrument (Microcal, Northampton, MA). Protein solutions were degassed, and the reference cell was filled with buffer. The cell volume was 0.51161 mL. Several authors have reported thermal unfolding of monoclonal antibodies with DSC, focusing on the midpoint temperature T_0_(19; 20). However, no data on the enthalpy of unfolding or on the effect of chemical denaturant are yet available. To the best of our knowledge, the present study is the first where thermal unfolding of an antibody is combined with chemical denaturation and analysed with respect to heat capacity and enthalpy.

In a DSC scan unfolding appears as an endothermic event that can be approximated as a Gaussian distribution curve. The temperature of the peak maximum is the midpoint temperature T_0_ and the area under the peak is the enthalpy change ΔH_cal_ of unfolding.

During the unfolding process, the heat capacity C_p_ of a protein goes through a maximum at the midpoint of the conformational transition. In addition, the post-transitional heat capacity is larger than that of the native protein by ΔC_p_.

The calorimetric unfolding enthalpy ΔH_cal_ is thus composed of the conformational enthalpy proper, 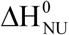 (often called van’t Hoff enthalpy ΔH_vH_), and the enthalpy increase 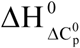, caused by the increased molar heat capacity 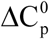 of the unfolded protein.

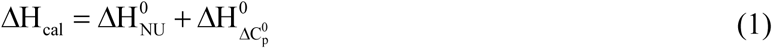

Antibodies exhibit a manifold of thermogram (19; 20). Due to the fact that monoclonal antibodies are complex, multi-domain protein, mAb unfolding is characterised by several independent unfolding domains (e.g. C_H_2, C_H_3, Fab). Proteins that contain multiple domains with different inherent stability require multiple Gaussians for empirical fitting of the thermograms. Protein denaturation is highly cooperative with many intermediates. For proteins/domains of 10-20 kDa size a simple 2-state model (all-or-none folding) is used as an approximation. When applied to antibodies (size ∼150 kDa) a single 2-state model leads to unacceptable results. Hence, the superposition of several 2-state models could be employed to generate an optimal fit. Empirical scaling factors must be used in addition and little insight into the molecular events is gained. In the present analysis we use a quantitative model based on the Zimm-Bragg theory (5). The model allows a truly cooperative analysis of antibody unfolding with deconvolution of individual domains.

It is common for therapeutic antibodies to aggregate and precipitate after unfolding. Especially at higher protein concentrations, after the main unfolding transition is passed, the heat capacity may drop sharply and then become negative. It is then not possible to obtain a correct post-unfolding baseline and, in turn, impossible to accurately evaluate the enthalpy of unfolding. The proper choice of the DSC baseline is an essential step in the quantitative analysis of DSC scans.

## Theory

### Thermal unfolding: Multi-state model (Zimm-Bragg theory)

We use N and U to denote the native and the unfolded conformation of the antibody, whereas n and u refer to a single amino acid residue.

We describe protein unfolding as a multi-state equilibrium between “native (n)” and “unfolded (u)” amino acid residues/peptide units (discussed in detail in a recent review(5)). A quantitative analysis is possible with the Zimm-Bragg theory (3; 4; 21). The essential parameters are the protein cooperativity σ and the equilibrium parameter q(T) of the native(n) → unfolded (u) transition of a single amino acid residue:

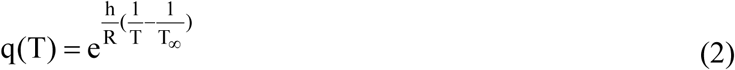

The enthalpy h of the n → u unfolding reaction is endothermic and is about 1.1 kcal/mol. h is an average value, comprising van-der-Waals interactions, electrostatic interactions and hydrogen bond formation (22; 23; 5).

The cooperativity parameter σ determines the steepness of the unfolding transition. A small σ corresponds to a high cooperativity. In the present study, the reference temperature T_∞_ is identical with the midpoint temperature T_0_.

The change in Gibbs free energy per amino acid for a temperature-induced unfolding in the interval T_ini_ ≤ T_0_ ≤ T_end_ is:

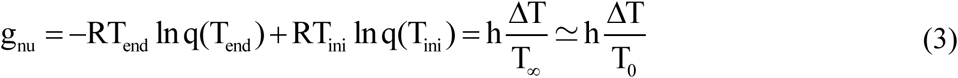

The free energy g_nu_ depends on the unfolding enthalpy h, the midpoint temperature T_0_, and the width of the unfolding transition ΔT = T_ini_-T_end_.

The central building block of the Zimm-Bragg theory is the partition function Z(T) = Z(σ,q(T)), which determines the statistical and thermodynamic properties of protein unfolding. Z(T) can be calculated with a matrix method (21):

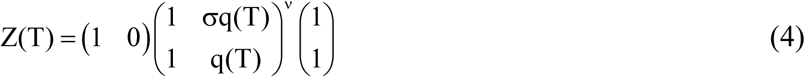

*v* is the number of amino acids involved in the unfolding reaction. q(T) is given by eq. (2). The fraction of native protein N, defined as Θ_N_ = Σn / *v*, is:

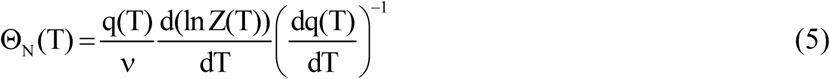

### Differential scanning calorimetry (DSC)

The transition from native protein (N) to the unfolded protein (U) is associated with an endothermic enthalpy ΔH_NU_(T) with the temperature-dependence:

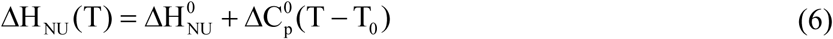

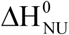 is the conformational enthalpy whereas the second term defines the contribution of the heat capacity increase 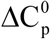. In the thermal unfolding experiment ΔH_NU_ (T) is convoluted with the extent of protein unfolding, Θ_U_(T) = 1-Θ_N_(T):

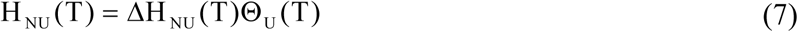

Differential scanning calorimetry measures the heat capacity:

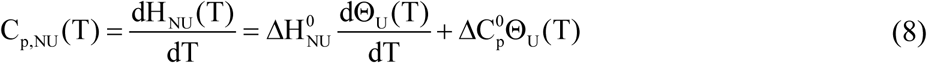

The enthalpy and entropy of unfolding are thus given by

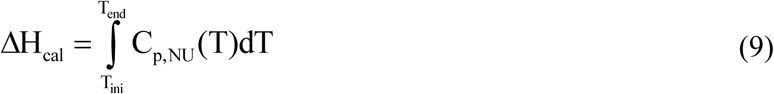

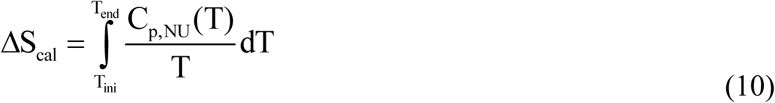

The contribution of the 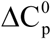 term is

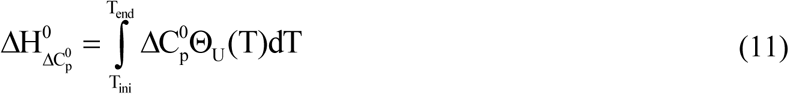

α-helix and β-sheet structures are usually assumed to require specific hydrogen bonds. Experimental studies on short alanine-based peptides contradict this classical view(24) as do free energy calculations using the CHARM potential function.(22; 23) Apparently, hydrogen bonds contribute little to α-helix/β-sheet stability since the major driving force favoring structure formation are enhanced van-der-Waals interactions and the hydrophobic effect.(22) Protein unfolding can thus be characterised by an *average* enthalpy h of approx. 1.1 kcal/mol per amino acid, independent of the specific protein conformation (5).

## Results

### Thermal unfolding (DSC) of mAb in guanidineHCl solution

Differential scanning calorimetry is the gold standard for thermodynamic analysis of protein unfolding, because thermodynamic data are directly obtained from the experiment. DSC measures the heat capacity C_p_(T) as a function of temperature and, by integration, the unfolding enthalpy H_NU_(T). The present mAb unfolding experiments in guanidineHCl solution were performed at pH 4.0, 6.2 and 8.0. Figure 1 shows the DSC scan of mAb in 1.0 M guanidineHCl at pH 6.2. The thermogram displays a low-temperature pre-transition and a high-temperature main transition, the general pattern of the present mAb unfolding experiments. Pre- and main transition are each characterised by a midpoint temperature T_0_ and an unfolding enthalpy ΔH_cal_. The number of amino acids involved in unfolding is *v* ≈ ΔH_cal_ / h. The averages of all measurements are *v* = 220 ± 50 for the pre-transition and 850 ± 30 for the main transition.

**Figure 1.**
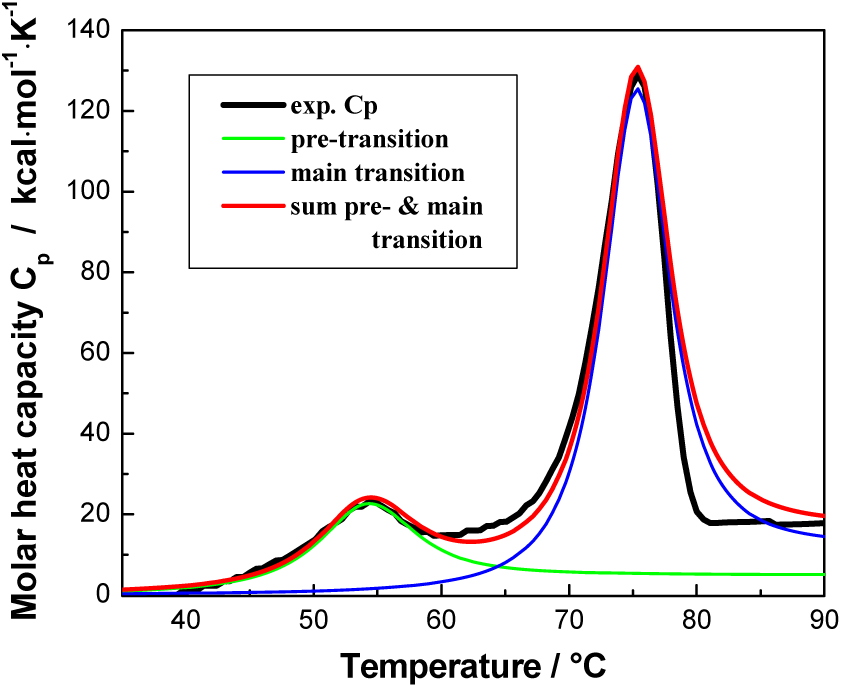
Thermal unfolding of mAb in 1.0 M guanidineHCl at pH 6.2, measured with differential scanning calorimetry (DSC). Molar heat capacity C_p_(T) as a function of temperature. Black noisy line: experimental result. Smooth lines: simulations with the multi-state Zimm-Bragg theory (green: pre-transition; blue: main transition; red: sum of pre- and main transition). Pre-transition parameters: T_0_ = 54 °C, ΔH_cal_ = 322 kcal/mol (v = ΔH_cal_/ h = 293); ΔC_p_ =5.02 kcal/mol·K, σ = 1.5×10^-4^. Main transition parameters: T_0_ = 75.4 °C, ΔH_cal_ = 976 kcal/mol (v= 887); ΔC_p_ =12.42 kcal/mol·K, σ = 7×10^-5^.

The multi-state Zimm-Bragg theory provides an almost perfect simulation of the DSC thermogram (Figure1 smooth red line) resulting from the superposition of pre- and main transition. The heat capacity of the unfolded protein is 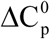 larger than that of the native mAb. Similar effects are well documented for thermograms of small proteins (25; 26). 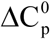 is caused by a restructuring of solvent molecules (25). The data in reference (25) suggest a linear relationship

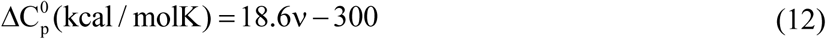

where *v* is the number of amino acid residues involved in unfolding. The heat capacity changes in Figure 1 are 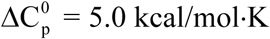 for the pre-transition and 12.4 kcal/mol·K for the main transition. Using equation (12) the numbers of amino acid residues are estimated as *v*∼280 for the pre-transition and 680 for the main transition, in broad agreement with the results derived from ΔH_cal_ with *v*∼290 and 890, respectively. Similar increases in the molar heat capacity of antibodies can be found in published DSC thermograms (e.g. (19; 27)).

Unfortunately, most DSC studies ignore the 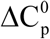 effect. The change in heat capacity between native and unfolded protein is eliminated by applying a sigmoid baseline. This choice of baseline results in a reduced unfolding enthalpy (e.g. (28)). The enthalpy of this truncated heat capacity peak is usually considered to represent the conformational enthalpy proper (also called “van’t Hoff enthalpy” in the 2-state model). However, “it is clear that in considering the energetic characteristics of protein unfolding one has to take into account all energy which is accumulated upon heating and not only the very substantial heat effect associated with gross conformational transitions, that is, all the excess heat effects must be integrated” (26).

In some mAb experiments, particularly at pH 4.0 and low guanidineHCl concentrations, the heat capacity drops sharply and becomes negative after the main unfolding transition. This can be explained by the formation of aggregates and perhaps precipitation after denaturation.

### Midpoint temperature T_0_ as a function of the guanidineHCl concentration

The midpoint of the unfolding transitions T_0_, defined by the C_p_ maximum, shifts linearly towards lower temperatures with increasing denaturant concentration (Figure 2).

**Figure 2.**
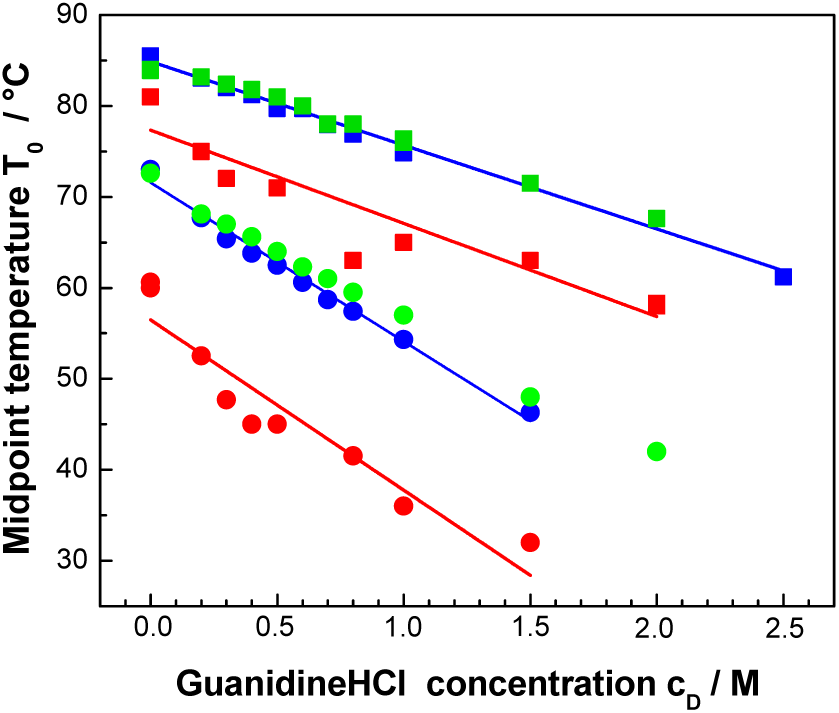
Midpoint temperature T_0_ as a function of denaturant concentration c_D_.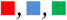 Main transitions at pH 4.0, 6.2 and 8.0, respectively. 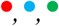 Pre-transitions at pH 4.0, 6.2, and 8.0, respectively.

DSC thermograms at pH 6.2 and 8.0 show almost identical transition temperatures at a given guanidineHCl concentration. At pH 4.0, the antibody is destabilized. The unfolding temperatures of pre- and main transitions are reduced by 15 °C and 7 °C, respectively.

Linear regression analysis of the data shown in Figure 2 yields for the pre-transition:

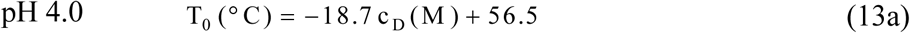

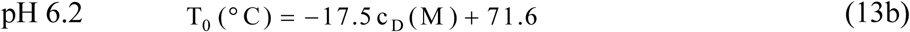

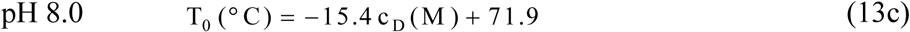

The guanidineHCL concentrations for mAb denaturation at 25 °C are predicted as 1.7 M (pH 4.0), 2.7 M (pH 6.2), and 3.0 M (pH 8.0).

The results for the main transition are:

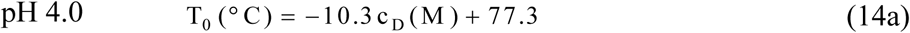

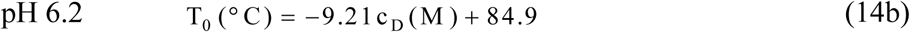

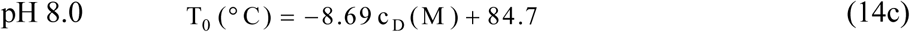

The guanidineHCL concentrations for denaturation of the mAb main transition at 25 °C are predicted as 5.1 M (pH 4.0), 6.5 M (pH 6.2), and 6.9 M (pH 8.0). The maximum solubility of guanidineHCl in water at room temperature is however only ∼ 6 M.

The pre-transition is twice as sensitive to guanidineHCl denaturation as the main transition.

### Unfolding enthalpy ΔH_cal_ as a function of guanidineHCL concentration

The calorimetric unfolding enthalpy, ΔH_cal_, decreases with increasing denaturant concentration c_D_ (Figure 3).

**Figure 3.**
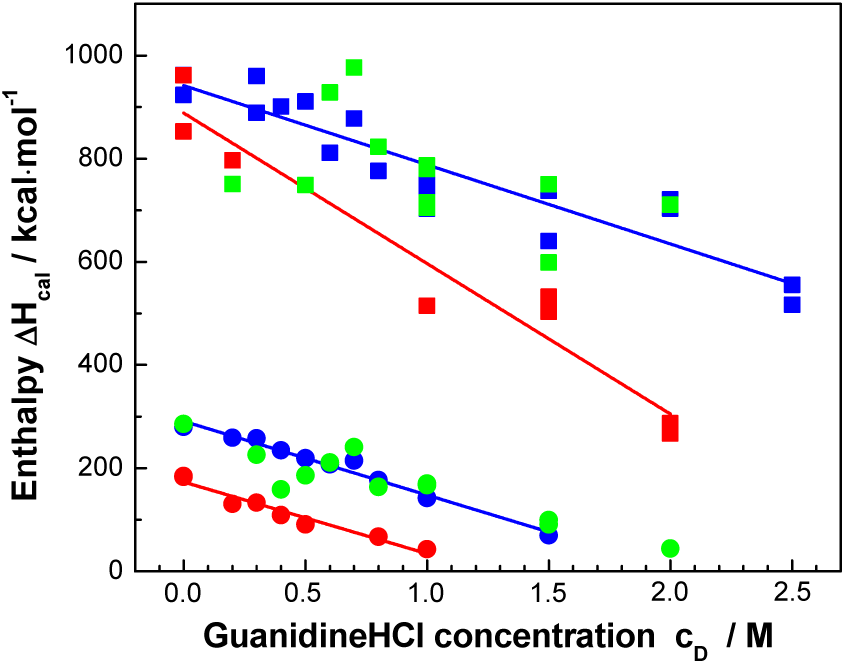
Unfolding enthalpy ΔH_cal_ as a function of denaturant concentration c_D_. 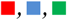 Main transitions at pH 4.0, 6.2 and 8.0, respectively. 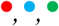 Pre-transitions at pH 4.0, 6.2, and 8.0, respectively.

Linear regression analysis yields for the pre-transition:

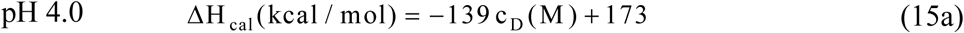

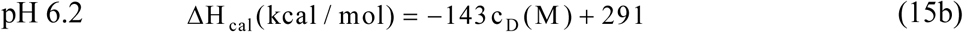

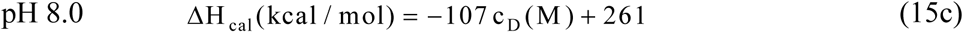

The number of amino acid residues involved in the unfolding transition can be estimated according to *v* = ΔH_cal_/h and are 157 (pH 4.0), 265 (pH 6.2), and 237 (pH 8.0) (Average: 220±50).

The results for the main transition are:

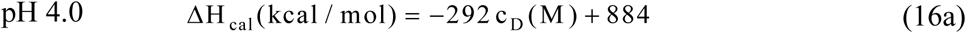

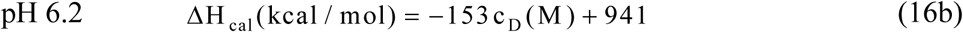

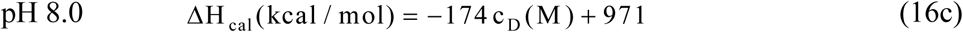

The number of amino acid residues is *v* = 808 (pH 4.0), 855 (pH 6.2), and 883 (pH 8.0).

(Average: 849 ± 30)

Antibody chains are divided into regions or domains consisting of around 110 amino acids. The pre-transition would thus represent the unfolding of about 2-3 domains, the main transition that of 8-9 domains. We tentatively assign the pre-transition to the reversible unfolding of C_H_2 domains and the main transition to Fab and C_H_3 domains.

### Unfolding enthalpy ΔH_cal_ as a function of midpoint temperature T_0_

The unfolding enthalpy ΔH_cal_ and the midpoint temperature T_0_ correlate linearly with the denaturation concentration c_D_. This predicts to a linear correlation between ΔH_cal_ and T_0_.

As shown in Figure 4, the enthalpies of pre- and main transitions cluster in narrow intervals. The slopes of the ΔH_cal_ versus T_0_ plots define a heat capacty 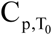. The average values are 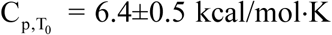 for the pre-transitions and 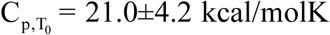 for the main transitions. The magnitude of the molar heat capacity 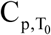 correlates with the number of amino acids *v* involved in the transition. In particular, 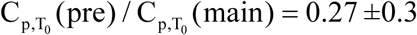 is identical within error to *v*(pre)/ *v*(main)= 0.30.

**Figure 4.**
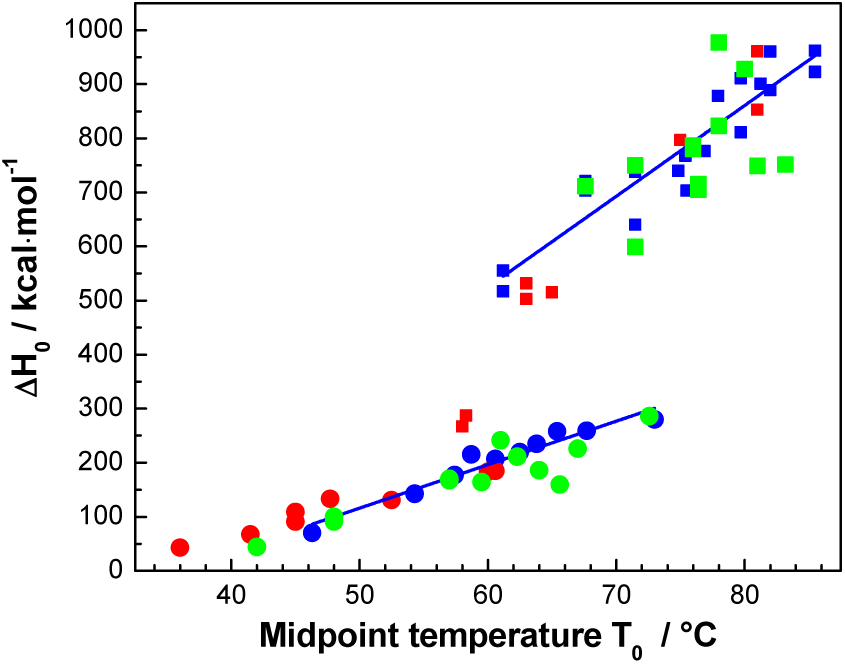
Unfolding enthalpy ΔH_cal_ as a function of midpoint temperature T_0_.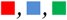 Main transitions at pH 4.0, 6.2 and 8.0, respectively. 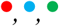 Pre-transitions at pH 4.0, 6.2, and 8.0, respectively.

### Cooperativity parameter σ

Figure 5 summarizes the cooperativity parameters for pre-transitions and main transitions. The cooperativity parameter σ increases slightly with increasing denaturant concentration, that is, the cooperativity of protein unfolding decreases. Figure 5 further demonstrates that σ increases from pH 8.0 (green symbols) over pH 6.2 (blue symbols) to pH 4.0 (red symbols). The σ parameter varies between 1.5×10^-5^ and 1.5 ×10^-4^ and is thus 10 to 100 times larger than σ of small proteins such as ubiquitin or lysozyme (cf. reference(5), Table 3).

**Figure 5.**
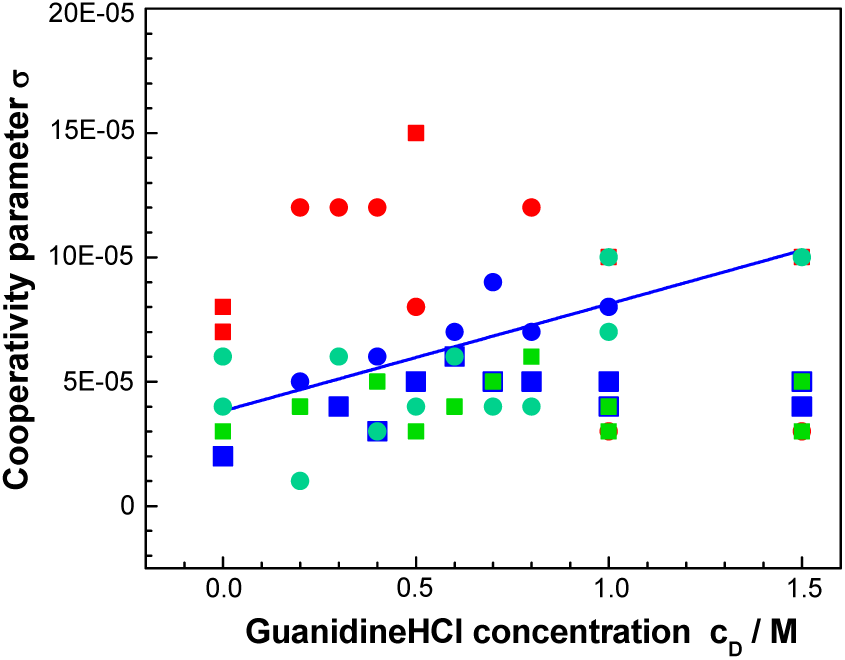
Cooperativity parameter σ as a function of guanidineHCl concentration c_D_.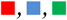 Main transitions at pH 4.0, 6.2 and 8.0, respectively. 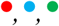 Pre-transitions at pH 4.0, 6.2, and 8.0, respectively.

The cooperativity parameter σ determines the average length <1> of a folded region according to 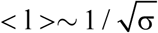. A cooperativity parameter σ = 10^-4^ thus predicts an average length of <1> = 100 amino acid residues. Several domains of length <1> will unfold independently and simultaneously upon heating mAb.

## Discussion

### Analysis of the DSC thermograms with the multi-state Zimm-Bragg theory

The 2-state model cannot fit the mAb pre- or main transition. In fact, it generally fails when applied to thermograms of large proteins. The unfolding of mAb is a multi-state transition with a large number of intermediates. In commercial instruments, an empirical curve-fitting software is applied without providing physical insight.

The multi-state Zimm-Bragg theory fits the mAb unfolding transition with 3 physical parameters: (i) the unfolding enthalpy h per amino acid residue, (ii) the cooperativity parameter σ and, (iii) the number *v* of amino acids residues involved in the transition. A small σ reflects a high protein cooperativity, which together with a large 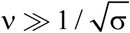 leads to a sharp DSC transition. In contrast, a large σ and/or a small *v* result in a broad transition with low cooperativity.

The Zimm-Bragg theory makes predictions about the average number <k> and average length <1> of segments that fold independently.(29) At the midpoint of the main transition of the thermogram shown in Figure 1 (T_0_= 75 °C) the Zimm-Bragg theory predicts <k> = 3.7 segments < 1 > ≃ = 100 amino acids. They are in dynamic equilibrium with interspaced unfolded regions.

As a general approximation the Zimm-Bragg theory predicts the average length of a folded segment as 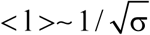.

The analysis of the unfolding enthalpies of small proteins led to an average unfolding enthalpy per amino acid of h = 0.9-1.3 kcal/mol (5). Hydrogen bonds have an unfolding enthalpy of 0.9 - 1.1 kcal/mol depending on the environment (4; 30). The present analysis used a constant value of h = 1.1 kcal/mol. As mentioned above, the number of amino acid residues participating in the unfolding reaction can be calculated as *v* = ΔH_cal_ / h. The exact value of *v* must not be known provided 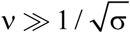. This condition is fulfilled for both pre- and main transitions, with cooperativity parameters of 1.5×10^-5^ ≤ σ ≤ 1.5×10^-4^. Cooperativity parameters of polypeptides and small proteins range between 10^-3^ for a 50 amino acid peptide and 5×10^-7^ for highly cooperative lysozyme(5). Most proteins of molecular weight 7-20 kDa have a cooperativity parameter of σ ∼ 10^-6^.

The Zimm-Bragg theory can be applied equally well to small and large proteins. It allows a comparison of molecular systems of different structure and size in terms of the parameters mentioned above.

### Antibody stability and unfolding temperature T_0_

The development of antibodies for therapeutic use has led to an increased effort in determining the factors influencing their stability. The stability of an antibody is dependent on different interactions such as van-der-Waals interactions, hydrophobic forces, hydrogen bonds, salt bridges, electrostatics, etc. In a DSC scan, these interactions are disrupted and the sum of all enthalpy changes is ΔH_cal_. Published DSC thermograms are, however, often analysed exclusively in terms of transition temperatures whereas ΔH_cal_ is ignored and not evaluated (19; 20; 27). Various attempts have been made to correlate the unfolding temperature T_0_ with structural characteristics of the antibody, assigning individual antibody domains to specific transition temperatures T_0_ (19; 20; 27). The associated enthalpy and entropy are however not considered even though these parameters are the essential factors in determining T_0_.

At the midpoint of the unfolding transition the Gibbs free energy is zero, ΔG (T_0_) = ΔH_cal_-T_0_ ΔS_cal_ = 0. It thus follows:

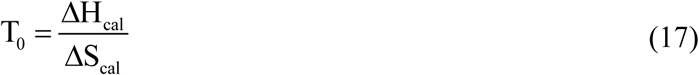

The transition temperature T_0_ is an indirect parameter, that is, it is the ratio of two thermodynamic quantities, the enthalpy ΔH_cal_ and the entropy ΔS_cal_. A small change in either enthalpy or entropy can lead to a significant change in T_0_. As an example, we compare the pre- and main transition of mAb unfolding at pH 6.2 in the absence of denaturant. The pre-transition is centered at 71.6 °C (eq. (13b)), the main transition at 84.9 °C (eq. (14b)). The corresponding enthalpies are ΔH_cal_ = 291 kcal/mol (eq. (15b)) and 941 kcal/mol (eq. (16b)). The entropies calculated with eq. (17) are ΔS_cal_= 0.84 kcal/mol·K and 2.63 kcal/mol·K, respectively. Next, the entropies are normalised with the number of amino acid residues involved. The entropy per amino acid residue is ΔS_cal_/*v* = 3.19 cal/mol·K for the pre-transition and 3.07 cal/mol·K for the main transition. The larger entropy of the pre-transition explains its lower melting temperature compared to the main transition if the enthalpy h is identical in both transition.

#### Unfolding enthalpy and the number of bound guanidineHCl molecules

GuanidineHCl decreases the unfolding enthalpy ΔH_cal_ of both the mAb pre- and main transition (Figure 3). In parallel, the transition temperature also decreases (Figure 2). According to T_0_=ΔH_cal_/ΔS_cal_ this is only possible if the entropy ΔS_cal_ changes less than the enthalpy ΔH_cal_.

The decrease of ΔH_cal_ and T_0_ with guanidineHCl is a general phenomenon which has been reported in DSC studies of small proteins such as lysozyme (11; 31; 32), ribonuclease (11), ubiquitin (33), and apolipoprotein A-1 (29). The present experiments are the first report for a large protein.

Guanidine is fully charged in the pH range of 4 to 8. A strong electrostatic interaction with charged peptide side chains was found (34). Recent X-ray studies of lysozyme also showed that guanidine binds to protein backbone and side chains and replaces water from the protein’s first solvent shell (13). GuanidineHCl binds to proteins with an exothermic binding enthalpy h_Gnd_ ≃ −2.63 kcal/mol (11) compensating, in part, the endothermic unfolding enthalpy ΔH_cal_. For mAb a concentration increase of Δc_D_ = 1 M reduces the enthalpy ΔH_cal_ of the pre-transition by δΔH_cal_ = −143 ± 5 kcal/molM (eq. (15b)). The number of bound guanidine molecules can thus be calculated as ΔN_Gnd_ = ΔδH_cal_/h_Gnd_ = (54 ± 2)/M. The corresponding results for the main transition are similar with ΔδH_cal_ = −(153 ± 10 kcal/molM (eq. (16b)) and ΔN_Gnd_ = (58 ± 5)/M.

Relevant for the unfolding reaction is the number of bound denaturant molecules after unfolding is complete. The midpoint concentrations 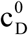 of the mAb pre- and main transition are predicted as 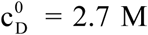 and 6.5 M, respectively (25 °C, pH 6.2). The pre-transition is completed at ∼3.5 M and N_Gnd_ = 190 guanidines are bound. The number of amino acid residues participating in the unfolding transition is N_aa_ = 265 (pH 6.2) leading to a stoichiometry of guanidineHCl/amino acid residues N_Gnd_/N_aa_ = 0.72. The main transition is completed at ∼7.3 M, resulting in N_Gnd_ = 424, N_aa_ = 855 (pH 6.2), and N_Gnd_/N_aa_ = 0.50. The pre-transition binds relatively more guanidineHCl molecules than the main transition.

The same analysis applied to published DSC data predicts for lysozyme (11) N_Gnd_ = 49±3, N_aa_ = 129; N_Gnd_/N_aa_ = 0.38; ribonuclease (11) N_Gnd_ = 49, N_aa_ =124, N_Gnd_/N_aa_ =0.4; ubiquitin (33) N_Gnd_ = 15, N_aa_ = 76, N_Gnd_/N_aa_ = 0.2; and apolipoprotein A-1 (29) N_Gnd_ = 50, N_aa_ = 110, N_Gnd_/N_aa_ = 0.45. Average N_Gnd_/N_aa_ = 0.36±0.09 (0.41±0.03 without ubiquitin).

#### Free energy of unfolding

The free energy g_nu_ of the n →u transition of a single residue depends on the width ΔT of the transition and the midpoint temperature T_0_ (eq. (3)). The width of pre- and main transition is ΔT ≃ 30 - 35 °C with about 95 % unfolded protein at the higher temperature. The free energy is thus g_nu_ ≃ 95 - 110 cal / mol, which is in agreement with results obtained previously with small proteins such as lysozyme or ubiquitin (5).

A completely different line of experiments supports the present results. The binding of amphipathic peptides/proteins to phospholipid membranes induces α-helix- or β-sheet structure. The Gibbs free energy change of the folding reaction was found experimentally and model-independent to be −140 to −400 cal/mol per amino acid (35-40). This result is of similar magnitude but of opposite sign than the mAb unfolding free energy g_NU_. The binding of amphipathic peptides to phospholipid *promotes* structure formation, and the free energy change is *negative*. In contrast, the binding of denaturants *disrupts* protein structure and the free energy change is *positive*. The two processes are of different sign but of equal magnitude.

## Concluding remarks

Thermal unfolding of a monoclonal antibody mAb displays two independent folding domains. A smaller domain (∼220 amino acids) centered at 72 °C and a larger domain (∼850 amino acids) with a midpoint temperature at 85 °C. Together the two domains account for ∼90% of the total amino acids of mAb. Unfolding is not a simple 2-state equilibrium between a fully folded and a fully unfolded domain (all-or-none equilibrium) but a complex reaction with many intermediates. The multi-state Zimm-Bragg theory provides an excellent description of the experimental data. The analysis of the DSC thermograms yields the unfolding enthalpy, the protein cooperative parameter σ, the number of residues participating in the unfolding, and the Gibbs free energy for the unfolding of a single amino acid residue. The theory predicts 10-12 segments of average length <1> ∼100, which fold independently and are in dynamic equilibrium. The addition of guanidineHCl up to 2.5 M has little influence on the protein cooperativity, but decreases drastically the unfolding enthalpy. The binding of guanidineHCl to the protein backbone and side chains is an exothermic reaction, which compensates in part the endothermic unfolding enthalpy. The decrease in the unfolding enthalpy yields the number of guanidineHCl molecules bound to each of the two domains. The stoichiometry guanidineHCl-to-amino acids is 0.72 for the small domain and 0.50 for the large domain. The small domain (C_H_2) is better accessible to the denaturant and thus easier to destabilize.

## Author Contributions

Antibody preparation and DSC measurments were performed by P.G., A.E., and M.B. Theoretical analysis were made by J.S.

## Acknowledgement

Work supported by the foundation “Stiftung zur Förderung der biologischen Forschung”.

## Abbreviations

mAb: monoclonal antibody
Gnd: guanidineHCl
DSC: differential scanning calorimetry
ITC: isothermal titration calorimetry
HPSEC: High Pressure Size Exclusion Chromatography
N: native protein
U: unfolded protein
n: native amino acid residue
u: unfolded amino acid residue
C_p_: heat capacity of the unfolding process
T_0_: temperature of the C_p_ maximum (midpoint temperature of N→U transition)
ΔH_cal_: calorimetric determined unfolding enthalpy of the transition
ΔH _vH_: van’t Hoff enthalpy
h: unfolding enthalpy per amino acid residue
σ: cooperativity parameter
q(T): equilibrium parameter for n → u transition
*ν*: number of amino acid residues participating in the unfolding reaction
g_nu_: free energy of the n → u transition of a single residue

